# Prodan-based solvatochromic probes for polarity imaging of organelles

**DOI:** 10.1101/2025.10.17.683001

**Authors:** Nathan Aknine, Paula Holban, Franziska Ragaller, Pablo Carravilla, Cenk O. Gurdap, Agit Cetinkaya, Erdinc Sezgin, Andrey S. Klymchenko

## Abstract

Solvatochromic probes provide microscopic, structural and functional information on their targeted cellular compartments. In this field, the challenge lies in designing probes that are both sufficiently sensitive to environment, and specific in their subcellular localization. Here, we design Prodan-based polarity probes targeting the following organelles: mitochondria, endoplasmic reticulum, Golgi apparatus, lysosomes and lipid droplets. The new probes provide robust organelle targeting, except for the mitochondrial probe, whose targeting ability is cell type dependent. Due to operating range of Prodan fluorophore in the UV-Blue region, our probes can be easily combined with other fluorescent tags in the visible range. Therefore, polarity of sub-cellular compartments can be studied together with additional fluorescent reporters. We used these probes to show the polarity of organelle membranes in healthy cells and under starvation condition where we observed organelle-specific polarity remodeling. These probes are important addition to the repertoire of smart cellular probes and will find critical use in understanding spatiotemporal regulation of cellular physiology.

## Introduction

Biomembranes are ubiquitous, micro-heterogeneous and complex structures, home to a wealth of biological activity. Biomembranes constitute not only the outer cell surface, but also delimit most intracellular organelle membranes, thus ensuring compartmentalization of cellular processes and specific functions, like signaling, active transport, trafficking and organelle-organelle interactions. Their activity is intimately linked to lipid order – a key biophysical parameter describing organization of membrane lipids.^1^ Two extreme cases of high and low lipid order in synthetic membranes can be defined: liquid ordered (Lo) and disordered (Ld). The membranes in Lo phase are tightly packed and highly ordered structures built of saturated lipids and cholesterol, such as sphingomyelin (SM), with a high cholesterol (Chol) content.^2^ Conversely, a membrane phase with low lipid order contains more unsaturated lipids, less arranged, with greater hydration, and is therefore more polar. Lipid packing in cellular membranes is more dynamic, assuming differential lipid order and forming membranes with wide range of biophysical properties and distinct physiological roles.^1, 3-5^

Fluorescent solvatochromic probes have demonstrated their potential as an analytical tool for deciphering the structural environment of living samples. These have the advantage of enabling direct observation with great sensitivity, being able to detect minute variations in the observed parameter. To shed light on lipid order in biomembranes, fluorescent probes sensitive to the environment are among the most popular tools.^6^ One should particularly stress three types of environment-sensitive probes for lipid order: solvatochromic dyes that sense polarity and fluidity,^6^ (2) molecular rotors that measure local viscosity^7^ and (3) fluorescent flippers that measure membrane tension.^8^

Prodan^9^ and its lipophilic analogue Laurdan^10^ are the first solvatochromic dyes proposed for studies of lipid organization, and they still remain popular because of their high sensitivity to polarity^11, 12^ and strong response to membrane domains of varying lipid order.^13-15^ Even though their absorption in UV and emission in blue green regions is generally considered as a limitation, this spectral range could be convenient for combination with dyes and fluorescent proteins operating in spectral ranges spanning from yellow to near-infrared regions. This allows multiplexing with many red/orange fluorescent proteins and dyes. To target Prodan specifically to the outer leaflet of the plasma membranes, we recently reported probe Pro12A, where the Prodan fluorophore was substituted with a lipid anchor group, composed of an anionic sulfonate group and lipophilic dodecyl chain.^16^ This probe enabled visualization of plasma membranes polarity in the presence of fluorescent proteins of interest, opening up new opportunities in biomembrane research.^17^

Targeting fluorescent dyes to organelles with chemical ligands is an attractive approach in fluorescence microscopy. This strategy enables organelles to be visualized without the need of cell fixation and immunolabeling.^18-20^ The chemical ligand exploits cellular biochemical mechanisms to localize to the organelle of interest. Thus, mitochondrial targeting is achieved by using hydrophobic cations, in particular based on triphenylphosphonium ions, which accumulate at the inner membrane of mitochondria in living cells through the mitochondrial membrane potential Δψ_m_.^21^ Lysosomes are targeted using a protonable moiety, usually weakly basic amines,^18^ which accumulate in these cellular compartments due to protonation at low pH. Lipid droplets are targeted utilizing the hydrophobicity of the organelle.^20, 22^ ER labelling strategies exist, which frequently include thiol-reactive groups such as alkyl chlorides or pentafluorophenyl esters.^23^ Finally, Golgi apparatus can be targeted using fatty acid derivatives, because it is responsible for processing lipids in the cells. Similar to endoplasmic reticulum, it is non-trivial to rationalize the design of fluorescent probes targeting the Golgi apparatus. Golgi probes are highly diverse, and their targeting mechanism is often misunderstood. However, one can assume that the similarity of probes targeting the Golgi with lipidic groups, usually processed by the Golgi results in their accumulation in this organelle. Classically, ceramide and myristoyl residues are good candidates for Golgi targeting.^18, 19, 24^

The targeting ligands can be attached to a fluorophore of interest with specific spectroscopic and sensing properties. They can be coupled to environment-sensitive fluorophores to visualize lipid organization in organelles.^25^ For example, fluorescent flippers were successfully targeted to mitochondria, endoplasmic reticulum, and lysosomes, allowing studies of membrane tension and lipid order in these organelles.^19^ Solvatochromic probes were particularly popular for targeting and sensing specific organelles, which includes mitochondria,^26, 27^ endoplasmic reticulum,^28^ Golgi apparatus,^29^ lysosomes,^30^ lipid droplets,^31^ etc. Recently, we showed that solvatochromic dye Nile Red can be targeted to different specific organelles using corresponding ligands, which enabled to study multiple organelles using the same fluorophore.^20^ This allowed comparison of lipid order in different organelles and decode organelle-specific response to external (mechanical and chemical) stress. Very recently, using similar strategy, Wong and Budin suggested to target Laurdan to specific organelles.^32^

In the present work, a new class of Prodan-based solvatochromic probes were synthesized, targeting five different organelles: mitochondria, lysosomes, endoplasmic reticulum, Golgi apparatus and lipid droplets (**Figure 1**). In contrast to the recent report on Laurdan derivatives, we used Prodan unit, which is more polar and thus ensure better solubility in water. The obtained new derivatives preserved sensitivity of Prodan unit to lipid order in model membranes. Moreover, taking advantage of blue operating range of Prodan, we performed co-localization studies with protein-based markers of organelles, which allowed us to evaluate the specificity of organelle targeting. These studies confirmed their specific targeting of organelles, although mitochondria targeting proved to be less reliable and dependent on the cell type. Finally, owing to Prodan solvatochromism, these probes reveled specific polarity profiles in the organelles of live cells under normal and starvation conditions.

**Figure 1.**
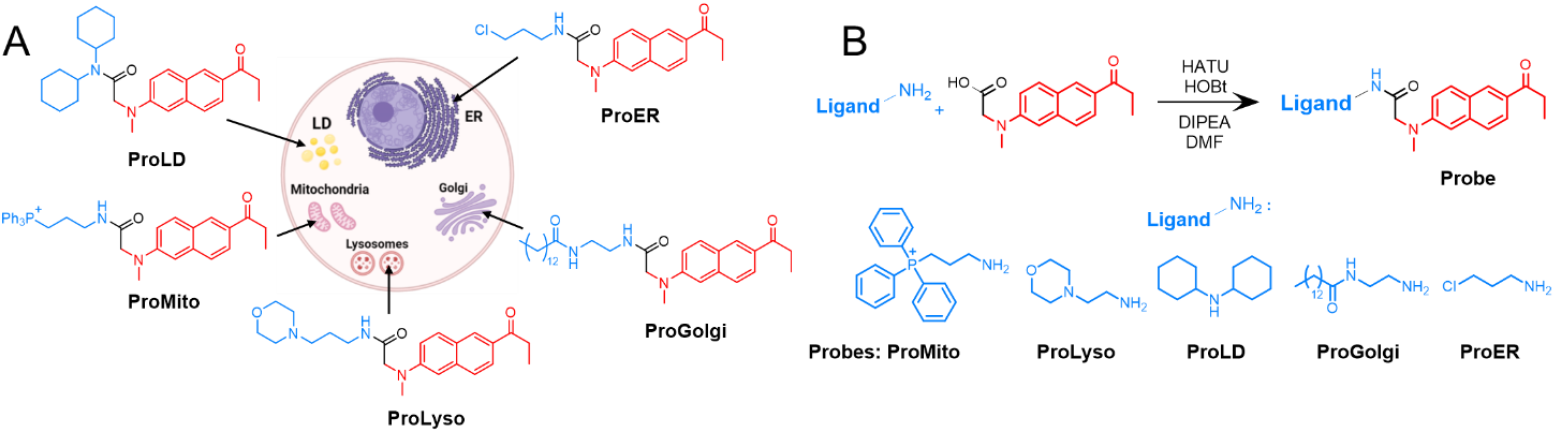
Design and synthesis of new probes. A) Prodan-based fluorescent solvatochromic probes targeting organelles: mitochondria (ProMito), lipid droplets (ProLD), endoplasmic reticulum (ProER), Golgi (ProGolgi), lysosomes (ProLyso). B) Synthesis route of Prodan-based organelle probes.

## Results and discussion

### Probe synthesis

The Prodan-based solvatochromic probes targeting organelles were obtained by conjugating a previously described Prodan fluorophore bearing a carboxy group Pro-CO_2_H with targeting moieties bearing an amino group (**Figure 1**).^16^ The coupling reaction was done using HATU agent in the presence of a base (diisopropylethylamine) and HOBt, followed by purification of the products by column chromatography. The products were obtained in moderate to good yields and confirmed by NMR and mass spectrometry.

### Probe characterization in solvents

After the synthesis, the probes were spectroscopically characterized in four solvents of varying polarity to assess their solvatochromism. The results were compared to Laurdan, which is an analogue of Prodan,^33^ bearing the same fluorophore, but a longer aliphatic chain, commonly used to study biological membranes.^10^ One can observe that functionalization of the fluorophore Prodan with organelle-targeting groups did not affect their polarity sensitivity: the emission band of the new probes shifted to the red in the more polar solvents similarly to that of the control dye Laurdan (**Figure 2**). On the other hand, there is a difference in fluorescence intensity between the probes in certain solvents: for example, ProMito, ProLyso and ProER are more emissive in the aqueous PBS buffer solution than the more hydrophobic derivatives, such as ProGolgi and Laurdan. These differences can be explained by the difference of solubility of these probes in water. It is clear that ProMito, ProLyso and ProER are more soluble in water than ProGolgi and Laurdan, bearing long alkyl chains, and therefore less likely to form non-emissive aggregates in PBS. These variations are quantified by the fluorescence quantum yield values in each solvent (Table S1). Their sensitivity to polarities confirms their potential use in ratiometric microscopy to assess local polarity profiles in organelles.

**Figure 2.**
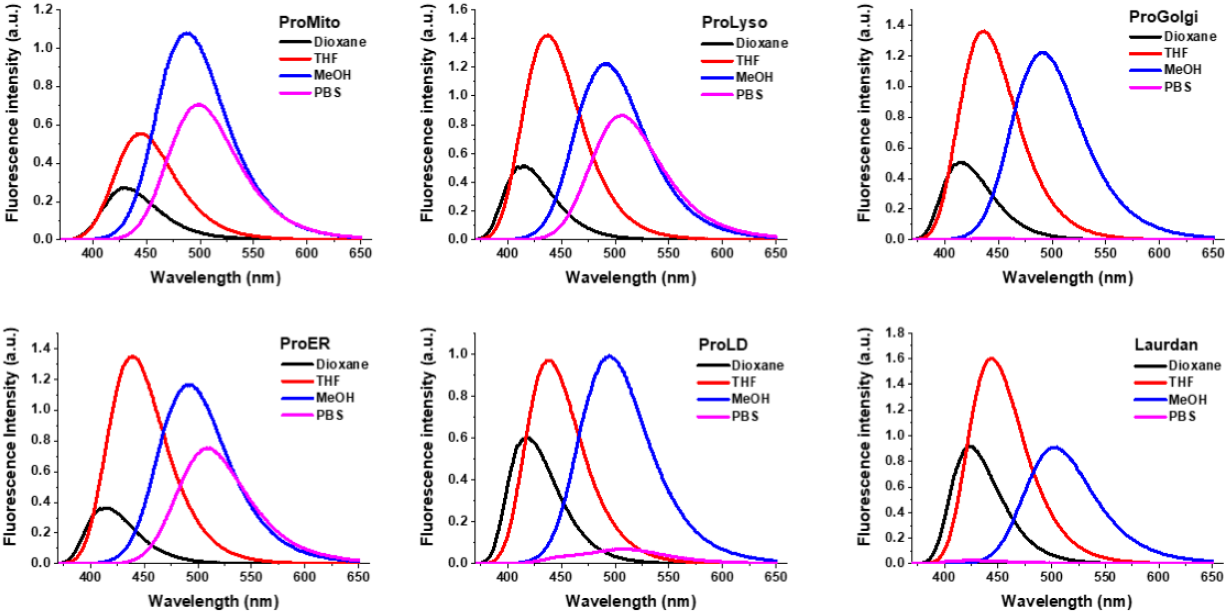
Emission spectra of solvatochromic probes in solvents: ProMito, ProLyso, ProGolgi, ProLD and ProER,as well as Laurdan, in solvents of increasing polarity: dioxane (black), THF (red), methanol (blue), and PBS (magenta). Probe concentration: 4 μM; λ (excitation) = 350 nm.

### Probe sensitivity to lipid composition

The sensitivity of the probes to lipid order was quantified by measuring their emission spectra in large unilamellar vesicles (LUVs), which are models of lipid bilayer membranes, presenting liquid disordered (Ld) phase (1,2-dioleoyl-sn-glycero-3-phosphocholine, DOPC, and its mixture with cholesterol, DOPC/Chol) and liquid ordered (Lo) phase (Sphingomyelin/Cholesterol, SM/Chol). LUVs were prepared by extrusion though a membrane with 100-nm pore in aqueous buffer, yielding vesicles of 130-nm diameter according to the dynamic light scattering (DLS). After incubation of the probes with these liposomes, their fluorescence spectra were acquired (**Figure 3**). Probes with the most polar structure, ProMito and ProLyso, showed fluorescence spectra close to that in the buffer (PBS), indicating that they failed to integrate into these lipid membranes due their high water solubility. More hydrophobic derivatives such as ProGolgi, ProER and ProLD incorporated into the vesicles, and their emission spectrum varied according to the nature of the lipids they contain. For ProGolgi, ProER and ProLD, the interpretation is clear: the higher the lipid order, the more blue-shifted the emission maximum was. These probes are therefore capable of distinguishing between different membrane phases of model membranes, similar to their parent analogues Prodan and Laurdan.^13-15^ Finally, ProER showed more complex behavior, because its emission in vesicles was strongly impacted by a significant contribution of emission from PBS. All the spectral characteristics for the Prodan-based organelle probes are summarized in Table S1.

**Figure 3.**
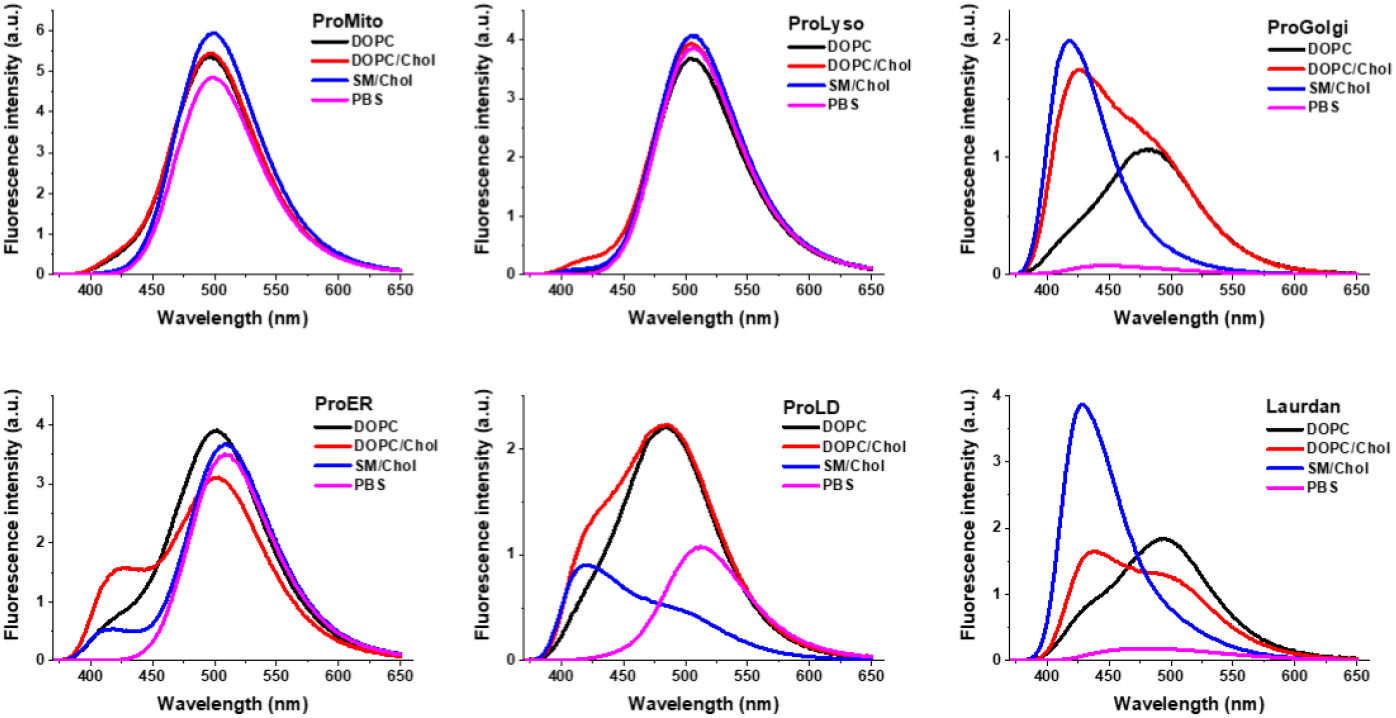
Emission spectra of ProMito, ProLyso, ProGolgi, ProER, ProLD, and Laurdan probes in LUVs of varying lipid composition: DOPC (black), DOPC/Chol (red), SM/Chol (blue), and PBS (magenta). Probe concentration: 400 μM. Total lipid concentration: 1 mM for each condition. λ (excitation) = 350 nm.

### Probe localization in live cells

These probes were then tested on different cell lines using cellular trackers (**Figure 4, Figure S1-3**). These results show an effective labeling for ProLyso, ProLD and Pro12A (our previously developed probe for PM) in all cell lines we tested (**Figure 4, Figure S1-3**). ProER, ProGolgi and ProMito localized well to the respective organelles in NRK cells, however, not in all cell lines. The main limitation with ProMito was the cell type dependent uptake of the probe. When the probe was uptaken to the interior of the cells effectively, the mitochondrial localization was also efficient. In HeLa and CHO cells, uptake of the probe was not efficient. ProER generally showed high colocalization with ER-tracker in all cell types. However, its colocalization with ER-specific protein KDEL was not as efficient. This is a general problem for ER trackers that they do not colocalize well with ER proteins.^34^ ProGolgi also showed cell type specific efficiency of localization. Similar to ER, ProGolgi colocalization with Golgi tracker was high in all cell types but not with Golgi protein Giantin in all cell types. In CHO cells, Giantin and ProGolgi colocalized perfectly while in NRK52E cells, colocalization was lower. Overall, while ProLyso, ProLD and Pro12A are effective in all cell lines we tested, ProMito, ProGolgi and ProER revealed two important limitations. First one, related to ProMito, suggest that targeting of some organelles is cell dependent. Second, localization of probes targeted by chemical ligands may differ from protein-based probes and this phenomenon also depends on the cell type.

**Figure 4.**
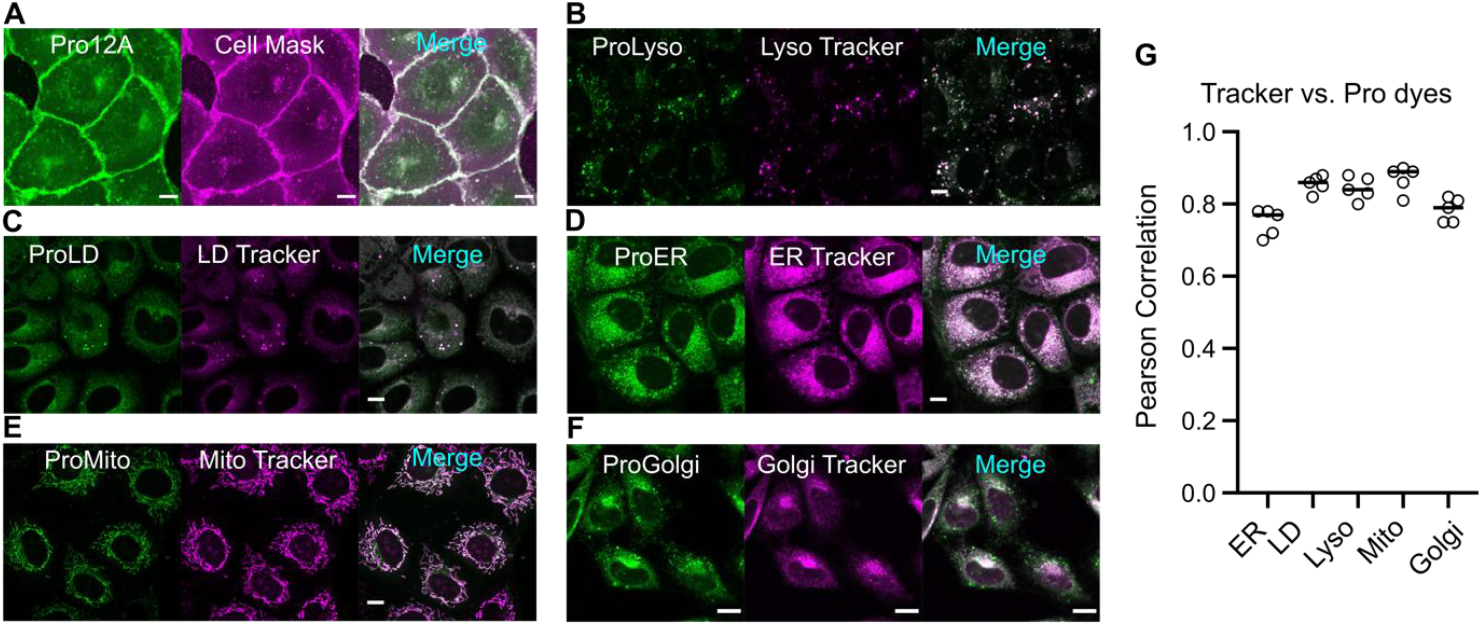
Confocal images of organelle trackers and Prodan-based organelle probes. A) Pro12A, B) ProLyso, C) ProLD D) ProER, E) ProMito, F) ProGolgi in NRK cells. (G) Pearson’s correlation coefficient for colocalization of Pro-probes vs. chemical trackers in NRK cells.

### Lipid packing of different organelles

Next, we tested whether we could compare the lipid order of different organelles in live cells. To explore the feasibility of such measurements, we first made model membranes of a fixed composition (bead supported lipid bilayers, BSLBs, composed of DOPC lipid) and measured the generalized polarization value (GP).^10, 13, 15^ The GP quantifies the red-shift of the probes according to differences in their immediate environment, with positive values representing a more ordered and negative values a more disordered environment. The GP of each probe depends on the lipid composition and order. However, the incorporation of the probes to the membranes was not uniform as shown in liposomes due to their varied polarity. Particularly ProMito showed poor incorporation and significantly different local polarity in BSLBs mostly dominated by noise, therefore it is not trivial to compare this probe with the others. ProLyso also poorly incorporated in BSLBs with very low signal, hence GP calculations from ProLyso was also noisy. Apart from the mitochondria probe, the rest of the probes showed similar GP values in the bead supported membranes composed of DOPC (**Figure 5a**), suggesting that these probes can be used to measure the differences between different organelles.

**Figure 5.**
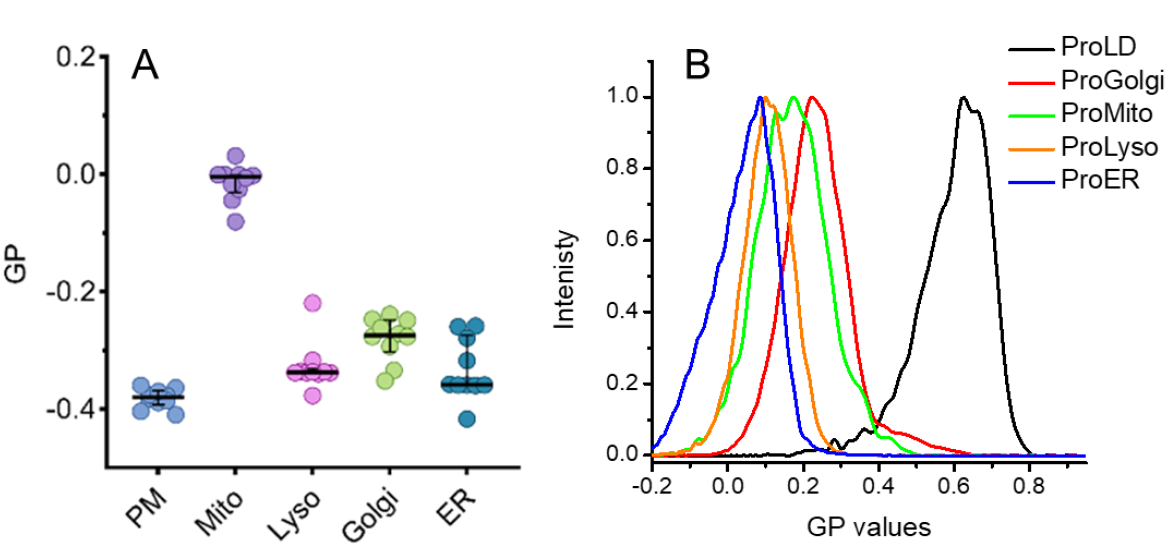
A) Lipid order analysis using the probes in BSLBs composed of DOPC. B) GP for the five Prodan-based solvatochromic probes in U87 cells.

To compare the polarity of the organelles, each probe was incubated with living cells and its emission was recorded in two distinct wavelength interval channels, namely: Channel A: [420-460] nm; and Channel B: [470-510] nm. These two channels correspond to the blue and red parts of Prodan emission band (**Figure 5**). Depending on the local polarity detected by the solvatochromic probe, its emission is expected to shift towards one or the other of the two channels. A good estimation of the environment polarity is therefore the intensity ratio between the two channels: this is the principle of ratiometric imaging. One B/A ratiometric image is produced from the two images of channels A and B, where pixel color reflects local polarity. In this case, the bluer pixels in the ratiometric image corresponds to less polar environment. All new probes were incubated with U87 cells, and the protocol described was applied in each case, generating ratiometric images (**Figure S4**). Moreover, from these data we extracted the distribution of GP values for each organelle targeting probe (**Figure 5b**). The latter suggested significant differences in the local polarity profile in the organelles (**Figure 5b**). The local polarity of the probes increased in the following order LDs < Golgi apparatus < Mitochondria < Lysosomes < ER. On the one hand, it can reflect differences in the local polarity and therefore order, suggesting that the most ordered membranes are located in the Golgi apparatus and the most disordered ones in the ER. The case of LDs is special, the observed lowest polarity in this case reflects the particulary low polarity of its lipid core, compared to biomembranes of other organelles that do not present an oil core. However, the obtained organelle-dependent polarity variation does not correlate well with the Nile Red-based organelle targeting probes.^20^ This indicates that the absolute values of polarity may not be compared for different probes, because they exhibit different localization in lipid membranes, in line with varied spectra observed in model membranes. Therefore, more detailed analysis with corresponding calibration should be done for each probe in order to compare lipid order in different organelles.^20^

### Differential response of organelles to stress

We next examined the influence of stress on the polarity of different organelles. To this end, we carried out serum-starvation for 24 h or 48 h and checked the GP value (membrane order) of plasma membrane, mitochondria and lysosomes (**Figure 6A-D**). Figure 6D depicts the changes in GP value upon starvation compared to non-starved cells. While the plasma membrane order increases upon starvation over time, no change can be observed for the mitochondria or lysosomes. This shows that different organelles are affected differently by cellular stress.

**Figure 6.**
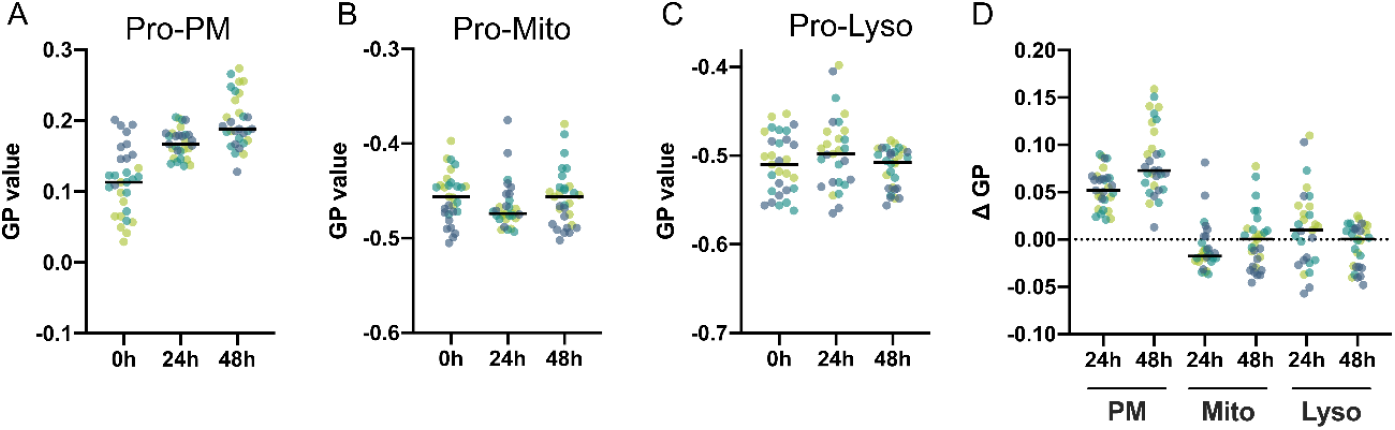
Ratiometric analysis of organelle remodeling during starvation. A) Pro-PM, B) Pro-Mito and C) Pro-Lyso for 24h and 48h of starvation. D) Remodeling calculated as changes in generalized polarization.

## Conclusion and perspectives

A new range of organelle-targeting solvatochromic probes based on Prodan has been developed, operating in the blue region of the visible spectrum. These probes are based on the affinity of certain targeting groups to the different organelles, namely mitochondria, lysosomes, Golgi apparatus and endoplasmic reticulum. Each probe was synthesized by peptide coupling between the fluorophore Pro-CO_2_H and corresponding targeting groups bearing an amino end. After conjugation by these groups, the new probes maintained their sensitivity to polarity and lipid order, characteristic of Prodan. This was demonstrated by their emission spectra in different solvents, highlighting positive solvatochromism, and in different model lipid membranes, highlighting sensitivity to lipid order.

Their targeting to organelles was assessed and confirmed on three cell types by colocalization experiments with known organelle trackers. Some new probes, like ProLyso and ProLD showed robust colocalization in all three cell lines. ProER and ProGolgi localized well with trackers based on chemical ligands, but their colocalization with protein markers depended on the cell type. This result shows that interpretation of localization of probes based on chemical and protein-based targeting should be taken with caution. Finally, ProMito showed that mitochondria targeting depends on the cell line.

Then, sensitivity of the Prodan fluorophore to polarity was used to evaluate the polarity profile of five different organelle membranes according to their membrane lipid order. To this end, ratiometric microscopy techniques were employed to quantify a reliable and reproducible polarity response. The results show that the lipid order of different organelles is not the same, although some differences might be probe dependent. It points out that care should be taken when different probes are used, highlighting the importance of probe calibration for data interpretation. The experiments on serum starvation revealed that this stress conditions could impact local polarity of plasma membranes but not lysosomes and mitochondria. These new probes could find applications in detecting biological events in organelles of living cells, in particular related to the cell stress. More generally, this new toolbox can be used in cell biology for multi-color imaging, thanks to Prodan’s emission spectrum, confined to the blue and green regions.

## Supporting information

Supplemental information for publication.

## Supporting Information

Supporting information is available.

## Acknowledgments

This work was supported by the Interdisciplinary Thematic Institute SysChem, via the IdEx Unistra (ANR-10-IDEX-0002), the CSC Graduate School (CSC-IGS ANR-17-EURE-0016) within the French Investments for the Future Program and Agence Nationale de la Recherche (ANR) AmpliSens ANR-21-CE42-0019-01. ES and his team have been supported by Swedish Research Council Grants (grant no. 2020-02682, 2024-02993 and 2024-00289), Wellcome Leap’s Dynamic Resilience Program (jointly funded by Temasek Trust), Karolinska Institutet (2024-03250; 2024-03341; 2022-00803; 2020-00997), Cancer Research KI (2024-03488), Human Frontier Science Program (RGP0025/2022), Longevity Impetus Grant from Norn Group, Hevolution Foundation and Rosenkranz Foundation. We thank the SciLifeLab Advanced Light Microscopy facility and National Microscopy Infrastructure (VR-RFI 2016-00968) for their support on imaging.

## Author contributions

NA and PH performed majority of the experiments, analyzed the results and NA and ASK prepared the first draft of the manuscript. ES, COG, FR, AC and PC performed some cellular experiments.

ASK and ES designed and supervised the project and analyzed some results. All authors contributed to manuscript writing and editing.

