## Supplemental information for publication. for "Prodan-based solvatochromic probes for polarity imaging of organelles"

### 1. General Methods

All starting materials for synthesis were purchased from Alfa Aesar, Sigma-Aldrich, TCI Europe, or ThermoFischer and used as received. MilliQ-water (Millipore) was used in all relevant experiments. NMR spectra were recorded on a Bruker Advance III 400 MHz or a Bruker Advance III 500 MHz spectrometer. NMR spectra were treated using MestRenova x64 software. Graphical data were edited using Origin software. High- resolution mass spectra were obtained using an Agilent 1200 ESI Q-ToF 6520 mass spectrometer. The size of lipidic particles was determined by Dynamic Light Scattering (DLS) with a Zetasizer ZSP (Malvern Instruments). ProCO2H was synthesized using a previously described procedure.[1]

### 2. Synthesis

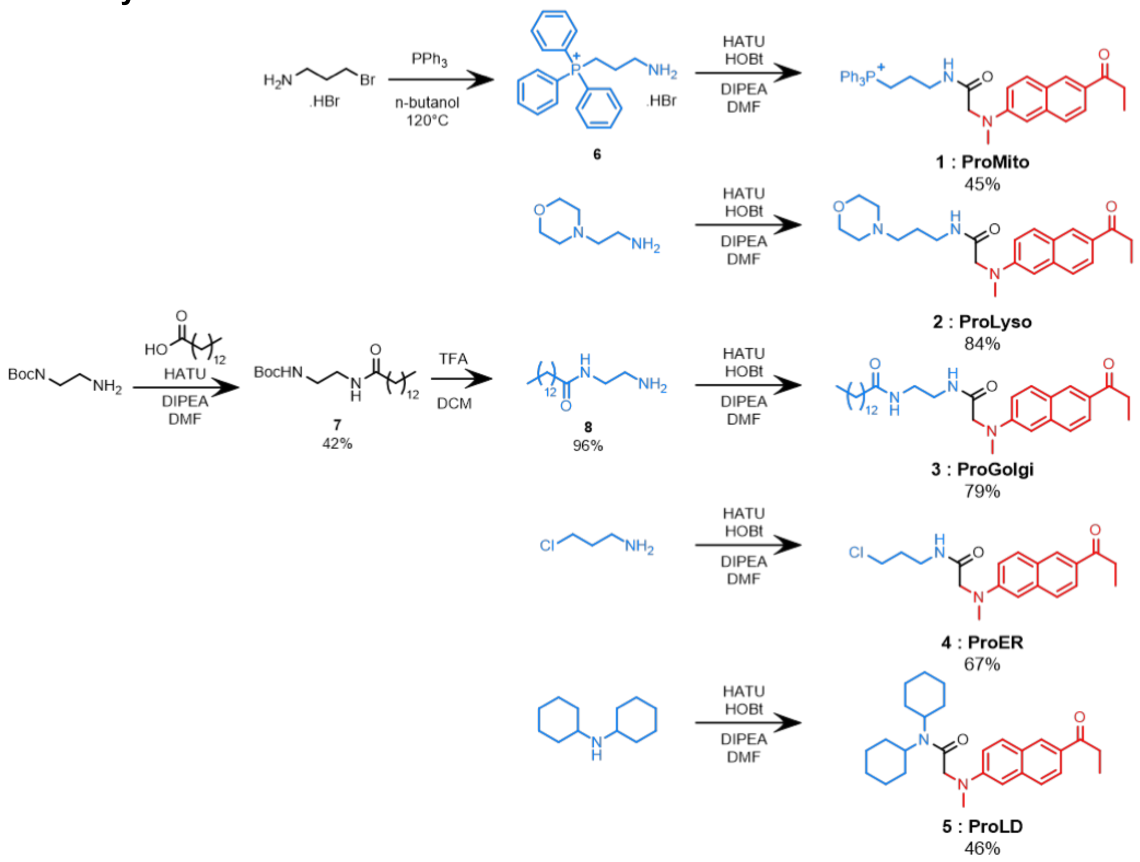


**Scheme S1.** Synthesis route of Prodan-based organelle probes: ProMito, ProLyso, ProGolgi, ProLD, et ProER.

**(3-(2-(methyl(6-propionylnaphthalen-2-yl)amino)acetamido)propyl)triphenylphosphonium bromide (ProMito) (1).** ProCO2H (60 mg, 221 μmol) was dissolved in 3 mL of dry DMF together with compound 6 (97 mg, 243 μmol) and DIPEA (154 μL, 884 μmol) under an Ar atmosphere. After 5 min, HATU (88.2 mg, 232 μmol) was added, and the mixture was stirred at RT for 16 h (controlled by TLC). The reaction was quenched with water, the solvent vas evaporated under vacuum and the crude product was purified by flash chromatography on silica SiO2 gel with gradual eluting with DCM/methanol from 100:0 to 92:8 v/v %. Yield: 65 mg (45%) as a yellow oil. $^1$H NMR (400 MHz, MeOD) δ 8.14 (d, J = 2.0 Hz, 1H), 7.76 – 7.67 (m, 3H), 7.65 – 7.47 (m, 8H), 7.47 – 7.31 (m, 7H), 7.07 (dd, J = 9.1, 2.6 Hz, 1H), 6.81 (d, J = 2.6 Hz, 1H), 4.00 (s, 2H), 3.34 – 3.26 (m, 2H), 3.20 (p, J = 1.7 Hz, 2H), 3.13 (s, 3H), 2.94 (q, J = 7.3 Hz, 2H), 1.65 (h, J = 7.1 Hz, 2H), 1.07 (t, J = 7.3 Hz, 3H). $^{13}$C NMR (101 MHz, MeOD) δ 201.34, 172.08, 149.03, 137.40, 134.94, 134.91, 133.24, 133.14, 130.72, 130.44, 130.21, 130.08, 129.71, 125.90, 125.52, 124.12, 118.41, 117.55, 115.88, 105.37, 56.43, 39.18, 38.76, 38.57, 37.49, 30.97, 29.28, 22.24, 19.47, 18.93, 7.62. MS (ESI), m/z: $[M + H]^+$ calcd for $C37H38N2O2P^+$, 572.2593; found, 572.2612.

**2-(methyl(6-propionylnaphthalen-2-yl)amino)-N-(3-morpholinopropyl)acetamide (ProLyso) (2).** ProCO2H (30 mg, 111 µmol) was dissolved in 3 mL of dry DMF together with HATU (44.1 mg, 117 µmol), HOBt (8.5 mg, 56 µmol), and (75.2 µL, 444 µmol) under an Ar atmosphere. After 5 min, 3-morpholinopropylamine (16.7 mg, 117 µmol) was added, and the mixture was stirred for 22 h (control by TLC) at r.t., under an Ar atmosphere. After the reaction, the solvent was evaporated in vacuo and the crude product was purified by column chromatography (SiO2; DCM/MeOH: 90/10). Yield: 37 mg (84%) as a yellow oil. $^1$H NMR (400 MHz, CDCl3): δ ppm 8.35 (d, J = 1.8 Hz, 1H) 7.97 (dd, J = 8.7, 1.8 Hz, 1H) 7.84 (d, J = 9.1 Hz, 1H) 7.68 (d, J = 8.7 Hz, 1H) 7.14 – 7.03 (m, 2H) 6.97 (d, J = 2.6 Hz, 1H) 3.49 – 3.44 (m, 4H) 3.39 (q, J = 6.3 Hz, 2H) 3.18 (s, 3H) 3.09 (q, J = 7.3 Hz, 2H) 2.32 – 2.16 (m, 6H) 1.65 (p, J = 6.6 Hz, 2H) 1.33 – 1.21 (m, 3H). $^{13}$C NMR (400 MHz, CDCl3 δ ppm 200.44 (Ccarbonyl), 169.86 (Camide), 149.03 (Car), 137.20 (Car), 131.67 (Car), 131.20 (Car), 129.49 (Car), 126.67 (Car), 126.27 (Car), 125.05 (Car), 116.26 (Car), 106.92 (Car), 66.75 (2Cal), 58.60 (Cal), 57.13 (Cal), 53.77 (2Cal), 40.09 (Cal), 38.60 (Cal), 31.66 (Cal), 25.44 (Cal), 8.63 (Cal). MS (ESI), m/z: [M + H]$^+$ calcd for C23H32N3O3, 398.2444; found, 398.2518.

**N-(2-(2-(methyl(6-propionylnaphthalen-2- yl)amino)acetamido)ethyl)tetradecanamide (ProGolgi) (3).** ProCO2H (20 mg, 74 µmol) was dissolved in 3 mL of dry DMF together with HATU (29.5 mg, 78 µmol), HOBt (5.7 mg, 37 µmol) and DIPEA (50.13 µL, 296 µmol) under an Ar atmosphere. After 5 min, a solution of N-(2-aminoethyl)tetradecanamide (20.9 mg, 78 µmol) and DIPEA (25 µL, 148 µmol) in 2 mL of DCM was added. The mixture was stirred for 28 h (control by TLC) under an Ar atmosphere. After the reaction, the solvent was evaporated in vacuo and the crude product was purified by column chromatography (SiO2; DCM/MeOH: 90/10). Yield: 30.5 mg (79%) as a pale yellow solid. $^1$H NMR (400 MHz, CDCl3): δ ppm 8.32 (d, J = 1.8 Hz, 1H) 7.93 (dd, J = 8.7, 1.8 Hz, 1H) 7.80 (d, J = 9.0 Hz, 1H) 7.64 (d, J = 8.7 Hz, 1H) 7.14 (t, J = 5.3 Hz, 1H) 7.05 (dd, J = 9.1, 2.6 Hz, 1H) 6.90 (d, J = 2.5 Hz, 1H) 5.98 (t, J = 5.4 Hz, 1H) 3.99 (s, 2H) 3.47 – 3.26 (m, 4H) 3.19 (s, 3H) 3.06 (q, J = 7.3 Hz, 2H) 1.91 – 1.82 (m, 2H) 1.43 – 1.04 (m, 25H) 0.87 (t, J = 6.8 Hz, 3H). $^{13}$C NMR (400 MHz, CDCl3) δ ppm 200.46 (Ccarbonyl), 174.49 (Ccarbonyl), 171.08 (Camide), 148.73 (Car), 137.28 (Car), 131.51 (Car), 131.15 (Car), 129.57 (Car), 126.64 (Car), 126.14 (Car), 125.00 (Car), 116.10 (Car), 106.60 (Car), 58.15 (Cal), 40.50 (Cal), 40.11 (Cal), 39.50 (Cal), 36.53 (Cal), 32.03 (Cal), 31.67 (Cal), 29.80 (Cal), 29.78 (Cal), 29.77 (Cal), 29.75 (Cal), 29.63 (Cal), 29.47 (Cal), 29.41 (Cal), 29.35 (Cal), 25.65 (Cal), 22.80 (Cal), 14.23 (Cal), 8.67 (Cal). MS (ESI), m/z: [M + H]$^+$ calcd for C32H50N3O3, 524,3853; found, 524.3900.

**N-(3-chloropropyl)-2-(methyl(6-propionylnaphthalen-2-yl)amino)acetamide (ProER) (4).** ProCO2H (30 mg, 111 µmol) was dissolved in 1 mL of dry DMF together with HATU (44.1 mg, 117 µmol), HOBt (8.5 mg, 56 µmol), and DIPEA (85.7 mg, 667 µL) under an Ar atmosphere. After 5 min, a solution of 3-chloropropylamine (15.1 mg, 56 µmol) in 1 mL of dry DMF was added, and the mixture was stirred for 24 h (control by TLC) under an Ar atmosphere, at r.t. After the reaction, the solvent was evaporated in vacuo and then the solid residue was dissolved in EtOAc, washed with water (x3) and brine, dried over Na2SO4, filtered and evaporated. The crude product was purified by column chromatography (SiO2; DCM/MeOH: 95/5). Yield: 25 mg (67%) as a yellow solid. $^1$H NMR (400 MHz, CDCl3): δ ppm 8.34 (s, 1H) 7.95 (dd, J = 8.7, 1.8 Hz, 1H) 7.83 (d, J = 9.1 Hz, 1H) 7.66 (d, J = 8.7 Hz, 1H) 7.07 (dd, J = 9.0, 2.6 Hz, 1H) 6.94 (d, J = 2.6 Hz, 1H) 6.68 (t, J = 6.2 Hz, 1H) 4.01 (s, 2H) 3.54 – 3.39 (m, 4H) 3.17 (s, 3H) 3.08 (q, J = 7.3 Hz, 2H) 2.00 – 1.92 (m, 2H) 1.25 (m, 3H). $^{13}$C NMR (400 MHz, CDCl3) δ ppm 200.54 (Ccarbonyl), 170.17 (Camide), 148.76 (Car), 137.22 (Car), 131.76 (Car), 131.32 (Car), 129.57 (Car), 126.72 (Car), 126.36 (Car), 125.11 (Car), 116.15 (Car), 106.97 (Car), 58.45 (Cal), 42.49 (Cal), 40.15 (Cal), 37.13 (Cal), 32.09 (Cal), 31.71 (Cal), 8.67 (Cal). MS (ESI), m/z: [M + H]$^+$ calcd for C19H24ClN2O2, 347.1527 ; found, 347.1582.

**N,N-dicyclohexyl-2-(methyl(6-propionylnaphthalen-2-yl)amino)acetamide (ProLD) (5).** Dicyclohexylamine (28.1 mg, 155.4 µmol) and DIPEA (50.1 µL, 296 µmol) were dissolved in 2 mL of dry DMF under an Ar atmosphere. Then, the mixture was stirred for 40 min at 50 °C. In a separate flask, ProCO2H (40 mg, 148 µmol) was dissolved in 1 mL of dry DMF together with HATU (58.9 mg, 155.4 µmol), HOBt (11.25 mg, 74 µmol), and DIPEA (100.3 µL, 592 µmol) under an Ar atmosphere.

After 10 min, the two solutions were mixed and the reaction mixture was stirred for 24 h at 45 °C (control by TLC). After the reaction, the solvent was evaporated in vacuo and the crude product was purified by column chromatography (SiO2; Heptane/EtOAc: 90/10). Yield: 28 mg (46%) as a pale yellow solid. $^{1}$H NMR (400 MHz, CDCl3): δ ppm 8.31 (d, J = 1.8 Hz, 1H) 7.91 (dd, J = 8.7, 1.8 Hz, 1H) 7.77 (d, J = 9.1 Hz, 1H) 7.59 (d, J = 8.7 Hz, 1H) 7.05 (dd, J = 9.1, 2.6 Hz, 1H) 6.83 (d, J = 2.5 Hz, 1H) 4.20 (s, 2H) 3.18 (s, 3H) 3.07 (q, J = 7.3 Hz, 2H) 2.42 (s, 2H) 1.92 – 1.42 (m, 12H) 1.37 – 1.01 (m, 11H). $^{13}$C NMR (400 MHz, CDCl3) δ ppm 200.61 (Ccarbonyl), 167.70 (Camide), 149.41 (Car), 137.75 (Car), 130.88 (Car), 130.70 (Car), 129.74 (Car), 126.42 (Car), 125.55 (Car), 124.60 (Car), 126.42 (Car), 115.92 (Car), 105.78 (Car), 57.42 (Cal),56.55 (Cal), 55.90 (Cal), 39.80 (Cal), 31.58 (2Cal), 30.16 (2Cal), 26.69 (2Cal), 26.13 (2Cal), 25.37 (2Cal), 8.78 (Cal). MS (ESI), m/z: [M + H]$^{+}$ calcd for C28H39N2O2, 435.3012 ; found, 435.3080.

**(3-aminopropyl)triphenylphosphonium bromide (6).** 3-bromopropylamine (835 mg, 3.81 mmol) was dissolved in n-butanol (20 mL) and triphenylphosphine (1g, 3.81 mmol) was added. The reaction was allowed to stir under reflux overnight. The reaction was cooled to RT and hexane (15 mL) was added. The suspension was stirred for 30 min and the precipitate was filtered and washed twice with hexane; The solid was triturated with a mixture of MTBE/hexane (50:50). The product was used in the next step without further purification. $^{1}$H NMR (400 MHz, MeOD) δ 7.87 – 7.62 (m, 15H), 3.61 – 3.49 (m, 2H), 3.12 (t, J = 7.7 Hz, 2H), 2.02 – 1.87 (m, 2H). $^{13}$C NMR (101 MHz, MeOD) δ 135.19, 135.16, 133.59, 133.49, 130.40, 130.27, 118.26, 117.40, 39.40, 39.18, 20.55, 20.52, 19.57, 19.03. MS (ESI), m/z: [M + H]$^{+}$ calcd for C21H23NP$^{+}$, 320.1568 ; found, 320.1580.

**tert-Butyl(2-tetradecanamidoethyl)carbamate (7).** Myristic acid (0.5 g, 2.18 mmol) was dissolved in 5 mL of dry DMF together with HATU (874 mg, 2.30 mmol), HOBt (148 mg, 1.09 mmol), and DIPEA (1.15 mL, 6.57 mmol) under an Ar atmosphere. After 5 min, a solution of N-Boc-ethylenediamine (368 mg, 2.30 mmol) in 5 mL of dry DMF was added, and the mixture was stirred for 24 h (control by TLC) at 50 °C. After the reaction, the solvent was evaporated in vacuo and the product was purified by column chromatography (SiO2, MeOH: 100). Yield: 340 mg (42%) as a pale yellowish solid. $^{1}$H NMR (400 MHz, CDCl3): δ ppm 6.12 (s, 1H) 4.90 (s, 1H) 3.40 – 3.31 (m, 2H) 3.31 – 3.22 (m, 2H) 2.16 (m, 2H) 1.61 (m, 4H) 1.44 (s, 9H) 1.25 (s, 18H) 0.87 (m, 3H).

**N-(2-aminoethyl)tetradecanamide (8).** Tert-Butyl(2-tetradecanamidoethyl)carbamate (150 mg, 405 µmol) was dissolved in 3 mL of dry DCM; then, 2 mL of TFA was added and the mixture was stirred for 2 h at room temperature. After the reaction, the solvent was evaporated in vacuo. In order to eliminate traces of TFA, 1 mL of methanol was added, followed by evaporation in vacuo (three times). Yield: 149 mg (96%) (in a form of TFA salt) as a yellowish oil. $^{1}$H NMR (400 MHz, CDCl3): δ ppm 7.24 (br s, 1H), 3.45−3.56 (m, 2H), 3.04−3.17 (m, 2H), 1.50−1.61 (m, 2H), 1.24 (s, 20H), 0.87 (t, J = 7.0 Hz, 3H).

### 3. Spectroscopy

Absorption and emission spectra were recorded on an Agilent Cary 5000 UV−vis−NIR spectrophotometer and an Edinburgh FS5 spectrofluorometer, correspondingly. The fluorescence quantum yields were determined using the formula of relative QYs:

$$\varphi_{f\ sample} = \frac{I_{sample}}{I_{ref}} * \frac{OD_{ref}}{OD_{sample}} * \frac{n^2{}_{sample}}{n^2{}_{ref}} * \varphi_{f\ ref}$$

With:
- I: integral of fluorescence spectrum
- OD: optical density (absorbance) at the excitation wavelength
- n: refractive index of the solvent

Fluorescence quantum yields were measured using as references:
- quinine sulfate in 0.05 M sulfuric acid (λex = 350 nm, QYref = 52%).[2]
- PK in DCM (λex = 420 nm, QYref = 99%).[3]

**Table S1.** Spectroscopic properties (absorption maximum, emission maximum and fluorescence quantum yield) of ProMito, ProLyso, ProGolgi, ProER, ProLD and Laurdan probes in organic solvents and model membrane vesicles (LUVs) of different lipid composition.

| Solvant/ liposome | paramètre | ProMito | ProLyso | Pro Golgi | ProER | ProLD | Laurdan |
|---|---|---|---|---|---|---|---|
| Dioxane | $\lambda_{max}$(abs) | 351 | 342 | 345 | 342 | 348 | 350 |
| | $\lambda_{max}$(em) | 429 | 414 | 414 | 415 | 420 | 424 |
| | QY (%) | 15 | 22 | 27 | 20 | 46 | 52 |
| THF | $\lambda_{max}$(abs) | 350 | 345 | 347 | 345 | 351 | 354 |
| | $\lambda_{max}$(em) | 444 | 437 | 437 | 440 | 436 | 444 |
| | QY (%) | 41 | 85 | 85 | 80 | 84 | 93 |
| MeOH | $\lambda_{max}$(abs) | 353 | 353 | 353 | 353 | 363 | 366 |
| | $\lambda_{max}$(em) | 487 | 492 | 489 | 491 | 494 | 505 |
| | QY (%) | 82 | 79 | 82 | 63 | 81 | 65 |
| PBS | $\lambda_{max}$(abs) | 365 | 363 | 321 | 363 | 377 | 346 |
| | $\lambda_{max}$(em) | 498 | 507 | 437 | 508 | 504 | 428 |
| | QY (%) | 69 | 60 | 2.2 | 57 | 23 | 3.9 |
| SM/Chol | $\lambda_{max}$(em) | 498 | 505 | 419 | 509 | 420 | 427 |
| DOPC/Chol | $\lambda_{max}$(em) | 498 | 504 | 427 | 503 | 484 | 437 |
| DOPC | $\lambda_{max}$(em) | 495 | 505 | 485 | 502 | 486 | 495 |

### 4. Preparation of Large Unilamellar Vesicles (LUVs)

All types of LUVs were prepared by the following procedure. A stock solution of corresponding lipid(s) in chloroform was placed into a round-neck flask, after which the solvent was evaporated in vacuo and phosphate buffered saline (PBS) was added. After all of the solid had been resuspended by vortexing, a suspension of multilamellar vesicles was extruded by using a Lipex Biomembranes extruder (Vancouver, Canada). The size of the filters was first 0.2 μm (7 passages) and thereafter 0.1 μm (10 passages). This generates monodisperse LUVs with a mean diameter of 0.13 μm, as measured with a Malvern Zetasizer ZSP (Malvern Instruments). The phospholipid/cholesterol molar ratio in the case of DOPC/Chol and SM/Chol was 1:0.9. For the preparation of SM/Chol LUVs, the temperature was systematically kept at 60 °C.

### 5. Cell lines, culture conditions, and treatments

**Culture.** U87 (ATCC HTB-14) cells were grown in Eagle's Minimum essential medium (EMEM, Gibco Invitrogen), supplemented with 10% fetal bovine serum (FBS, Lonza), 2 mM L-glutamine (Gibco-Invitrogen), 1% non- essential amino acid solution (Gibco-Invitrogen) and sodium pyruvate 1 mmol/L at 37 °C in a humidified 5% CO2 atmosphere. Cells were seeded onto a chambered coverglass (IBIDI) at a density of 5 × $10^4$ cells/well 24 h before the microscopy measurement.

**Staining.** Cell medium was removed from the well and the cells were washed with PBS. The PBS solution was removed and a dye solution in OptiMEM was incubated on cells:

- 45 min at 37°C for mitochondria probes
- 30 min at 37°C for lysosome, Golgi, and ER probes
- 10 min at RT for LD probes

After the incubation, the cells were imaged without further washing.

### 6. Fluorescence microscopy

**Confocal microscopy.** Confocal imaging of cells was performed on a Leica TCS SP8 confocal microscope with a HCX PL APO 63×/ 1.40 OIL CS2 objective and two 12-bit photomultipliers. The excitation light was provided by lasers of 405 and 638 nm. The fluorescence from the channels was detected at the spectral ranges depending the experiment (described in figure captions). All the parameters at each channel were left constant; the illumination power was adjusted to achieve a good signal for each probe. When the performances of the probes were compared, all instrumental conditions were fixed. All the images were processed using ImageJ software.

**Colocalization.** Colocalization analysis was performed using the plugin Colocalization Finder (developed by P. Carl, University of Strasbourg) under ImageJ. From two images as an input, the plugin generates a merged image and a Pearson's correlation coefficient that fluctuates between -1 (exclusion), 0 (random), and 1 (perfect colocalization).

For colocalization experiments, NRK-52E and HeLa cells were cultured in DMEM medium supplemented with 10% fetal bovine serum. CHO cells were grown in DMEM/F12 supplemented with 10% fetal bovine serum. Cells were seeded 24 hours before imaging in 8-well Ibidi µ-slides. The following day cells were labelled with Pro-dyes by adding them directly in growth medium. The DMSO concentration was kept at 0.2% (v:v). Pro-dye concentration was 1 µM Pro-PM, 2.5 µM Pro-Mito, 1 µM Pro-Lyso, 2 µM Pro-Golgi and 2 µM Pro-ER. Organelle marker labelling was performed simultaneously by adding the markers 5 min after adding Pro-dyes. Marker concentrations were those recommended by manufacturers, namely 400 nM MemBright 590 (Idylle), 10 nM MitoTracker Orange CMTMRos (ThermoFisher Scientific), 50 nM LysoTracker Red DND-99 (ThermoFisher Scientific), 240 µM Golgi-Tracker Red (GLPBIO) and 1 µM ER-Tracker Red (BODIPY™ TR Glibenclamide). Cells were then incubated for 20 min at 37 °C 5% CO2, except for plasma membrane labeling, which was imaged immediately after labelling to reduce dye internalization. After incubation with dyes and trackers, cells were washed twice with Leibovitz's L-15. For fluorescence protein-targeted labelling, cells were seeded 48 h before imaging and transfected 24 h prior with 50 ng ER-mScarlet-I or mNeonGreen-Giantin using Lipofectamine 3000. ER-mScarlet-I (Addgene plasmid # 137805; http://n2t.net/addgene:137805; RRID:Addgene_137805) and mNeonGreen-Giantin (Addgene plasmid # 98880; http://n2t.net/addgene:98880; RRID:Addgene_98880) were a gift from Dorus Gadella.
Images were acquired using a LSM 780 confocal microscope (Zeiss) equipped with a Zeiss C-APOCHROMAT water immersion objective lens (40x /1.2). Excitation/detection was performed in line-steps for each channel, i.e. dyes were not excited simultaneously. Two-line averaging was performed. Pro-dyes and organelle trackers were excited with 405 nm and 561 nm laser lines, respectively. mNeonGreen was excited with a 488 nm laser and mScarlet-I with a 561 nm laser. Emission was collected at 405-510 nm for Pro-dyes, and 570-691 nm for organelle trackers or mScarlet-I. In experiments with mNeonGreen, detection windows were 405-474 and 504-625 nm. The pinhole size was kept to 1.0 airy units. Images were analyzed using Fiji.

**Ratiometric imaging.** The ratiometric images were generated by using the plugin RatioloJ (developed by R. Vauchelles, University of Strasbourg) under ImageJ that divides the image of one channel by the other (emission windows of channels are described in figure captions). For each pixel, a pseudocolor scale was used for coding the ratio, while the intensity was defined by the integral intensity recorded for both channels at the corresponding pixel.

**Cell starvation.** For starvation experiments, 3x104 NRK cells per well were seeded in µ-Slides (18 well glass bottom, ibidi) and serum-starved (0% FBS) for 0h, 24h or 48h. Before imaging the cells were incubated with either 3.5 µM of Pro-Mito or 2 µM Pro-Lyso in DMEM for 20 min at 37 °C and washed with phenol red- and serum-free L15 medium before imaging. To investigate the plasma membrane the cells were incubated with 0.8 µM of Pro12A in phenol red- and serum-free L15 medium for 5 min and the dye solution was not removed before imaging. For confocal spectral imaging of the cells, a Zeiss LSM 780 confocal microscope with a 32-channel array of gallium arsenide phosphide (GaAsP) detectors was utilized. The probes were excited at 405nm, and the emitted fluorescence was collected in approximately 9 nm wavelength intervals between 423 and 601 nm (20 channels) simultaneously. The images were thresholded to exclude background noise and the general polarization value (GP) was subsequently calculated as $GP=(I_B-I_R)/(I_B+I_R)$, with $I_B$ and $I_R$ as the fluorescence signal intensities at the blue- or red-shifted emission wavelengths, respectively. The GP can range between +1 and −1, with positive values representing a more ordered environment and vice versa. The intensities in the channel at 423 nm (λLo) and 494 nm (λLd) were used for GP calculation.

**Coated beads production**. 10 µL of 25 mg/mL DOPC lipid solution was dried under nitrogen flow to evaporate chloroform. Dry film was hydrated with 1 mL of SLB buffer (150 mM NaCl, 10 mM HEPES, pH 7.4). Later, it was vortexed for a few minutes and tip sonication for 10 mins (power 3, duty cycle 40%) was applied. 45 uL of liposome solution was added to 5 uL beads with the final volume of 500 uL with SLB buffer. Bath sonication at room temperature was applied for 30 minutes. The resulting beads coated with lipid bilayers were washed three times with SLB buffer again. Beads were incubated with probes at a final concentration of 300 nM for 10 mins. They were washed once with SLB buffer, placed into an 18-well glass bottom ibidi chamber for imaging. Zeiss LSM 780 confocal microscope with 32-channel detectors was used as previously described.

### Supplementary Figures

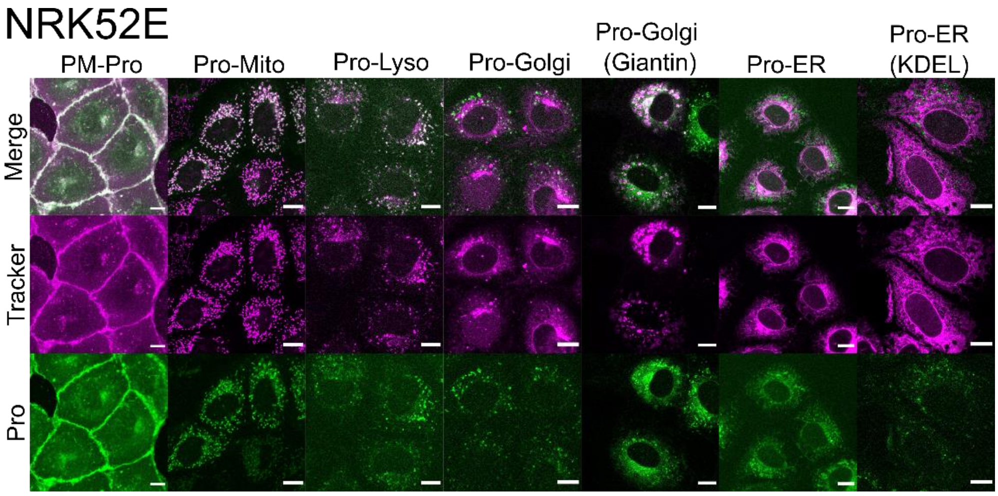


**Figure S1.** Colocalization experiments for all new probes in NRK-52E cells with corresponding trackers, using conforcal microscopy.

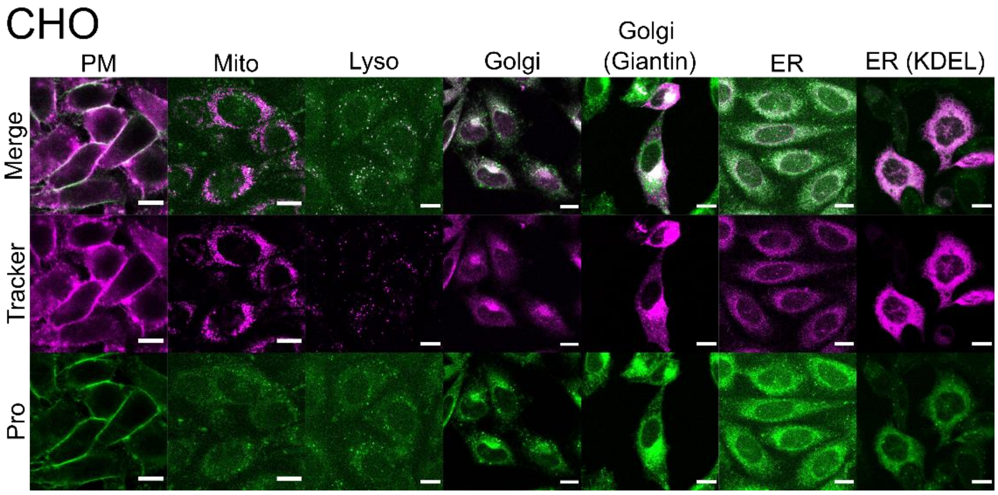


**Figure S2.** Colocalization experiments for all new probes in CHO cells with corresponding trackers, using conforcal microscopy.

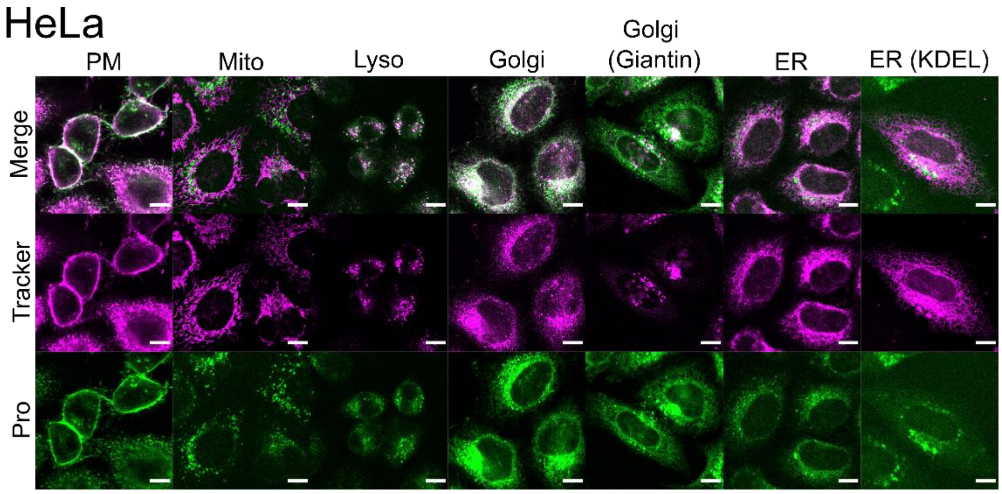


**Figure S3.** Colocalization experiments for all new probes in HeLa cells with corresponding trackers, using conforcal microscopy.

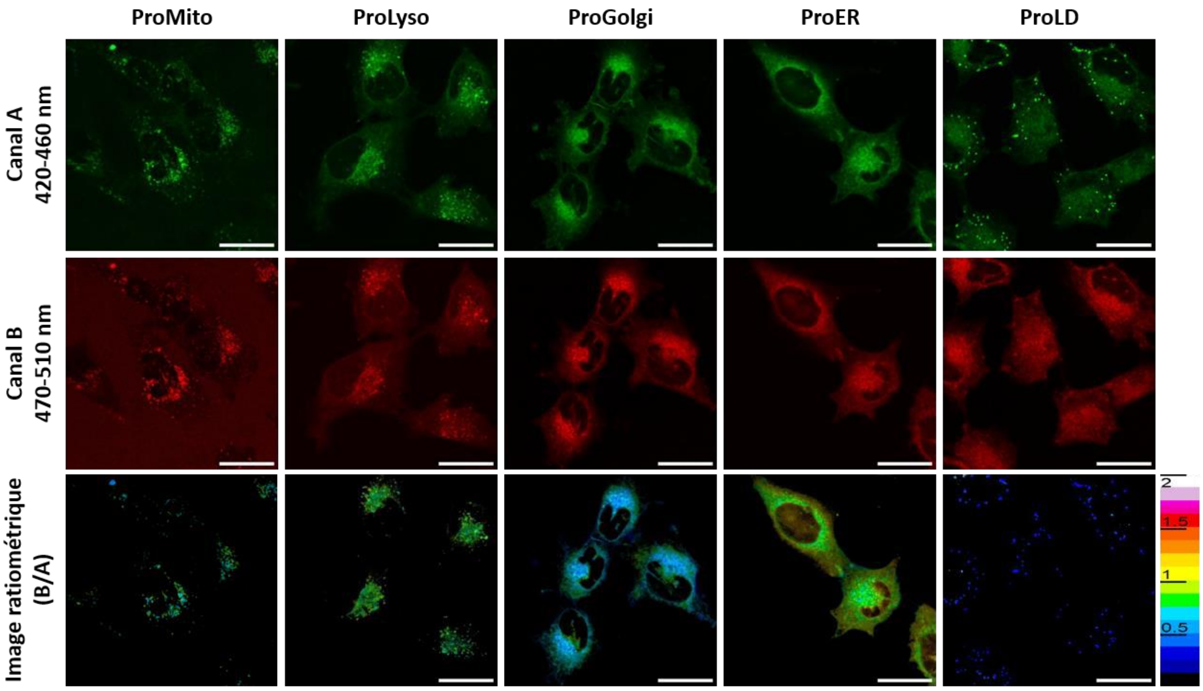


**Figure S4.** Confocal microscopy images of U87 cells labeled with ProMito, ProLyso, ProGolgi, ProER, and ProLD probes in two wavelength channels (red and green); channel A (green) = [420-460] nm and channel B (red) = [470-510] nm. Ratiometric images (B/A) were generated using RatioloJ on ImageJ. λ (excitation)= 405 nm. Scale bar: 30 µm.
